# StereoMapper: Clarifying Metabolite Identity Through Stereochemically Aware Relationship Assignment

**DOI:** 10.64898/2025.12.09.693222

**Authors:** Jack McGoldrick, Marco Pagni, Saleh Alwer, Joanne Cooney, Natalia Makosa, Anne Niknejad, Sébastien Moretti, Jadzia Murphy, Filippo Martinelli, Alan Bridge, Ines Thiele, Ronan M.T. Fleming

## Abstract

Inaccuracies in metabolite cross-mapping frequently introduce errors into genome-scale metabolic models (GEMs). A key challenge lies in distinguishing truly identical metabolites from closely related structural variants, such as stereoisomers, across biochemical databases. StereoMapper addresses this issue by establishing stereochemically aware relationships that clarify metabolite identity. It operates entirely on structural information, using molecular structures to infer equivalence and define relationships. StereoMapper was implemented using RDKit (2025.3.3) and Open Babel-wheel (3.1.1.22) which provides the core functionality of the Open Babel software, and benchmarked across four curated control datasets to evaluate performance in relationship classification (enantiomer, diastereomer, stereo-resolution pairs, and protomer). The pipeline was subsequently applied to 1.3 million molecular structures from multiple metabolic databases to generate equivalent isomeric sets (groups of structures identified as equivalent by StereoMapper). These sets were compared against the human subset of MetaNetX, a context-aware reference, to assess concordance and investigate disagreements. On curated positive control sets containing only positive examples across multiple relationship classes, StereoMapper achieved mean precision, recall, and F1-scores of 92.6, 98.0, and 95.3, respectively; following optimisation, these improved to 98.9, 98.2, and 98.5. Application to the 1.3 million structures produced over 339,000 stereochemically defined relationships between molecular structures. Comparison with the human subset of MetaNetX assignments showed an overall grouping concordance of 93.5%. The remaining 6.5% of differences were mainly associated with protomer relationships (52.3%) and stereo-resolution pairs (37.4%), while 21.6% involved complex or ambiguous cases.

**Scientific contribution statement:** StereoMapper provides a stereochemically aware pipeline for metabolite relationship assignment that shows high performance across multiple benchmark datasets, resolving key ambiguities in metabolite identity and providing a complementary approach to existing context-aware frameworks. Its integration will enhance reaction-level cross-mapping and improve the accuracy of future GEM reconstructions.

## 1 Introduction

Genome-scale metabolic models (GEMs) are computational frameworks that represent the complete metabolic network of an organism, linking genes to reactions and metabolites [1]. They enable prediction of metabolic phenotypes and have diverse applications, including disease biomarker identification, drug discovery in complex diseases such as cancer, and elucidating host–microbiome interactions [2, 3, 4]. Given their broad utility, GEMs have attracted increasing attention across systems biology, medicine, and bioengineering. Developing a GEM is a laborious and time-intensive process that typically involves multiple researchers with detailed domain knowledge of the target organism [5]. This effort relies heavily on specialised biochemical databases that provide metabolite, reaction, and genetic information critical for accurate reconstruction. Prominent examples include the Virtual Metabolic Human (VMH), Chemical Entities of Biological Interest (ChEBI), the Human Metabolome Database (HMDB), and Rhea [6–9]. Each of these resources focuses on different aspects of biochemical knowledge: VMH supports microbial and human reconstructions such as Recon3D [10], ChEBI provides a curated ontology of chemical entities; HMDB offers detailed metabolite and clinical data; and Rhea delivers expertly curated biochemical reactions linked to ChEBI identifiers. However, because each database has its own scope and data structure, integration and synchronisation of these resources are essential to capture all relevant information for model construction.

Significant challenges remain in harmonising data across biochemical resources, particularly in distinguishing truly equivalent metabolites from closely related structural variants such as stereoisomers or protonation forms. Several established cross-mapping frameworks support this process, including MetaNetX and BridgeDB [11, 12]. MetaNetX was originally developed to cross-map metabolites from the biochemically, genetically, and genomically (BiGG) models knowledge base [13] to other structure-containing databases, at a time when openly available metabolite structures were limited. It adopts a context-aware strategy in which equivalence is inferred primarily through co-occurrence in identical reactions across multiple databases, supported by structural information within its reconciliation workflow, and is widely used within model reconstruction pipelines. Through this workflow, MetaNetX can group structural variants of a metabolite under a single identifier, leading to ambiguity in metabolite identity. BridgeDB, by contrast, provides an infrastructure layer for identifier translation by integrating mappings from multiple source databases and enabling transitive links across them. Because it relies on these upstream resources, any incorrect or inconsistent cross-references present at source may be carried through within BridgeDB. Both the reaction-based merging of structural variants in MetaNetX and the carry-over of inconsistent identifiers through BridgeDB can enter GEMs and lead to inappropriate cross-mappings, potentially introducing incorrect reactions, disrupting mass balance, or generating topological dead ends that affect model feasibility. [14–16].

Although context-aware reconciliation, as implemented in MetaNetX, incorporates structural information alongside reaction context, the combined use of these signals may provide limited resolution of fine-grained structural differences. Approaches that evaluate molecular structure directly therefore offer a complementary perspective, helping to distinguish structural variants that may be treated as equivalent within context-aware frameworks. These approaches have become increasingly feasible as high-quality metabolite structures and openly accessible cheminformatics toolkits have become more widely available. This now enables systematic evaluation of stereo-chemical variation, protonation state differences, and other subtle structural features that are not readily resolved through reaction co-occurrence alone. A structure-focused layer can therefore complement existing context-driven mappings by adding isomer-level detail, providing a more comprehensive basis for metabolite reconciliation that balances biochemical relevance with structural precision.

To complement existing context-aware reconciliation frameworks, we developed StereoMapper, a structure-driven tool that provides unambiguous assignment of structural relationships between metabolites. It enables clear differentiation between truly equivalent metabolites and related but distinct variants, such as stereoisomers or protonation forms. StereoMapper integrates established cheminformatics frameworks, including RDKit, Open Babel, and InChI, to systematically analyse and classify molecular relationships [17–19]. By defining explicit structure-based connections, it adds an additional layer of precision to existing context-aware mappings, thereby enhancing the clarity and reliability of metabolite cross-referencing across biochemical databases.

## 2 Methods

StereoMapper was developed following the series of steps illustrated in Figure 1, which are described in detail in the sections that follow. This section also outlines the procedures for data collection used in benchmarking and comparison against a context-aware framework, as well as the design and execution of the experimental analyses. The methods are organised to reflect the sequential stages of the StereoMapper development and evaluation process: Data Collection (Section 2.1), StereoMapper Workflow (Section 2.2), and Benchmarking Experiments (Section 2.3). StereoMapper was implemented with RDKit (2025.3.3), InChI software (1.07), and Open Babel-wheel (3.1.1.22) which provides the core functionality of the Open Babel package [18].

**Figure 1:**
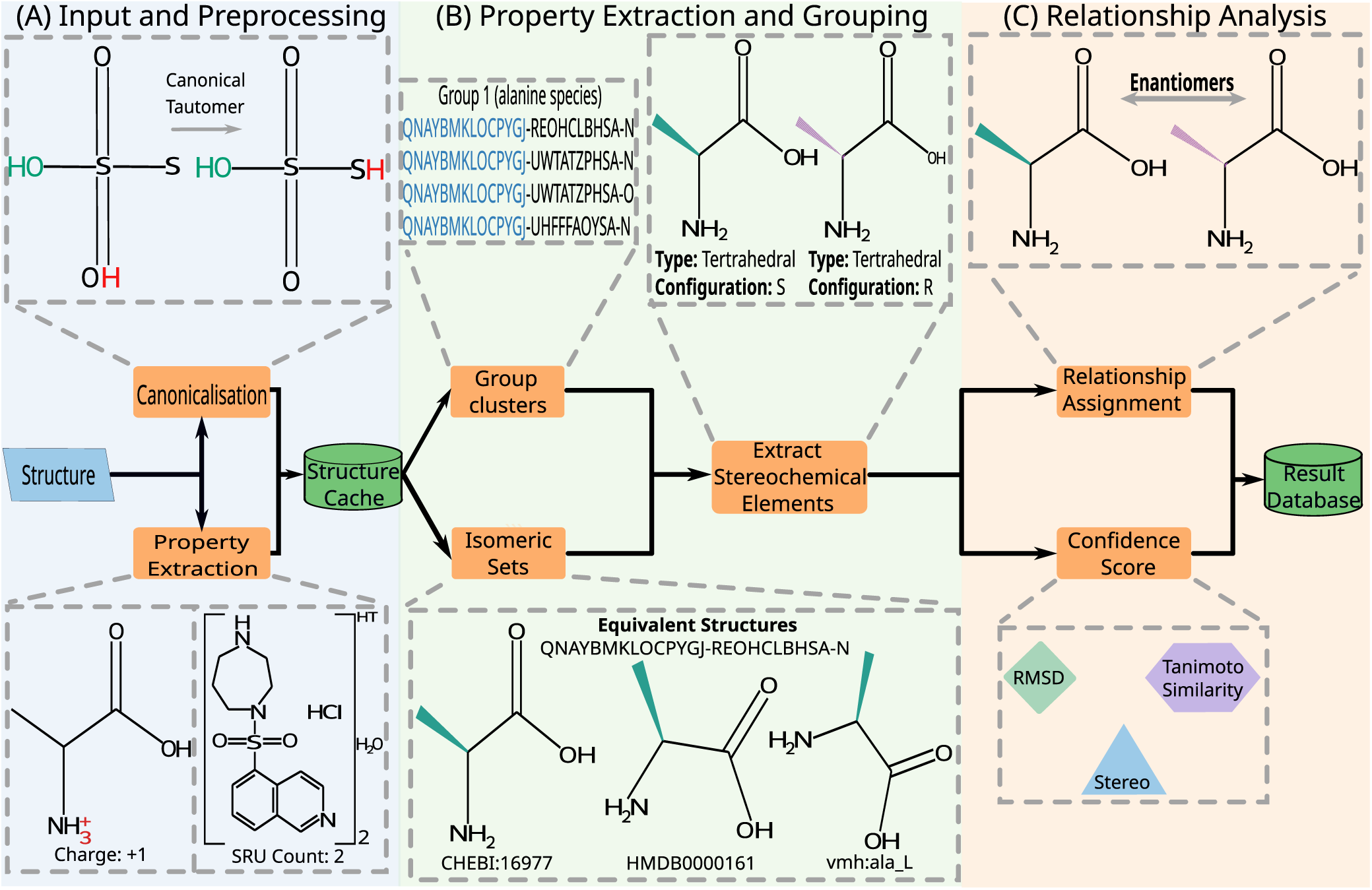
Overview of the StereoMapper pipeline. Each shaded region corresponds to a major processing stage, as indicated by the titles within each section: (A) Input and preprocessing (blue), (B) Property extraction and grouping (green), and (C) Relationship analysis and output generation (orange). Grey dashed lines indicate associations between visual elements and their corresponding processes. The diagram provides a high-level overview of the sequential workflow implemented in StereoMapper.

### 2.1 Data Collection

Data collection was performed in two phases to warrant experimental validation of StereoMapper: (i) generation of control datasets for benchmarking relationship assignments, and (ii) extraction of molecular structures from different biochemical databases used for application of the StereoMapper pipeline.

#### 2.1.1 Assembly of Benchmarking Control Datasets

Control datasets were constructed to evaluate the accuracy and precision of StereoMapper’s stereochemical relationship assignments. Four relationship classes were considered: (i) *enantiomers* are stereoisomers that are non-superimposable mirror images of one another and differ in configuration at every stereogenic centre, (ii) *diastereomers*are stereoisomers that are not mirror images and differ at one or more, but not all, stereogenic properties, including tetrahedral and cis/trans configurations, (iii) *protomers* are structures with identical structural connectivity and spatial configuration, but differing in protonation state, and (iv) *stereo–resolution pairs* are structure pairs where one structure is a more stereochemically resolved form of the other. Within the scope of StereoMapper, two structures are considered distinct variants if they possess identical atomic connectivity but differ in spatial configuration or protonation state, thereby warranting assignment to one of the relationship classes defined above.

For enantiomeric pairs, the ChEBI database was used as a reference benchmark, as it provides manually curated structural relationships, including explicit “is enantiomer of CHEBI:X” annotations. These data were downloaded as a flat file, from which enantiomeric pairs were obtained and, their corresponding structures were extracted. This process yielded approximately 1,300 enantiomeric pairs. A similar approach was used to retrieve protomeric pairs. ChEBI includes the relationships “is conjugate acid of CHEBI:X” and “is conjugate base of CHEBI:X”, which describe pairs sharing identical connectivity but differing in protonation state, aligning with the definition of protomers used in StereoMapper. To avoid duplication, only one of these relationships was used, resulting in over 8000 initial pairs. A structural diversity filter using molecular fingerprints generated with RDKit [17] and 2D Tanimoto coefficient [20] (threshold = 0.7) was used to remove structurally redundant pairs. In this context, a Tanimoto similarity of 1.0 corresponds to identical fingerprints, whereas 0 indicates no overlap, and pairs with similarity *≥* 0.7 were treated as redundant. The final control dataset contained approximately 3,300 protomeric pairs. Stereo resolution pairs were derived analogously by retrieving parent-child relationships from MetaNetX, which represent the same structural relationship but are expressed using alternative terminology. To maintain consistency with other datasets, only relationships between ChEBI structures were retained, yielding over 5000 pairs.

Since diastereomer relationships are not manually curated in ChEBI or other databases, we developed a multistep approach combining InChIKey grouping, stereochemical feature analysis, and manual validation. InChIKeys were generated for all ChEBI structures using RDKit’s InChI interface [21], and candidates were grouped by identical first and third blocks but differing second blocks, which encode various types of isomeric information, including stereochemistry [19]. Because the initial grouping of candidate diastereomers produced many false positives, exhaustive manual inspection was impractical. We therefore refined the dataset using RDKit-based stereochemical analysis to retain only pairs consistent with diastereomerism. This approach, which mirrors logic used within the StereoMapper pipeline, substantially reduced the number of pairs requiring review. As with the protomeric dataset, the 2D Tanimoto coefficient (threshold = 0.7) was applied to reduce redundancy. The resulting diastereomer control set contained over 3,097 pairs. Manual validation of all pairs would have been time-consuming and prone to fatigue-related errors. Instead, a stratified random sample of 315 pairs was manually reviewed, confirming 312 (99%) as true diastereomers (95% CI: 97.2–99.7%), corresponding to an estimated overall error rate of ∼1%. Manual review involved inspection of SMILES [22], InChI and InChIKey strings, alongside visualisation of molecular structures to confirm adherence to the definition of diastereomers [23]. In total, the curated control dataset comprised 1,362 enantiomeric, 3,097 diastereomeric, 3,365 protomeric, and 5,068 stereo-resolution pairs, providing a comprehensive benchmark for evaluating StereoMapper performance across relationship classes.

#### 2.1.2 Structural data extraction and alignment with context-based frameworks

Metabolite structural information used to test StereoMapper was collected from several biochemical databases, including the VMH, ChEBI, HMDB, KEGG, SwissLipids, LIPID MAPS, SABIO-RK, and ModelSEED [6, 7, 8, 24, 25, 26, 27, 28]. Structural data were obtained through a combination of direct downloads (molfile or SDF format), API requests, and flat-file access. To ensure inclusion of only structures retrieved directly from the source databases, identifiers from each were aligned with those in the human subset of the context-aware framework MetaNetX. Here, the human subset refers to the portion of MetaNetX metabolites known to be present in the human body. A single MetaNetX identifier can encompass multiple distinct variants of a metabolite originating from different databases, and all metabolites sharing the same MetaNetX identifier belong to the same cluster. Each StereoMapper cluster groups isomerically equivalent molecules, sharing atomic connectivity, stereochemical configuration, and protonation state under a common StereoMapper identifier (see Section 2.2.2 for details). For alignment, all metabolite identifiers within MetaNetX were retrieved, and each distinct member was cross-referenced against data obtained directly from the corresponding databases. Only members present in both MetaNetX and their respective source databases were retained, ensuring the benchmarking set contained structures consistently represented across both sources. Importantly, StereoMapper excludes generic structures from its analysis, as these are not supported in the current version of the pipeline. Generic structures include those that contain an R-group or wildcard atoms, indicating that they represent structural scaffolds rather than fully defined molecular entities. This process yielded a dataset of approximately 1.3 million structures. In parallel, each member and its associated MetaNetX identifier were recorded and linked to the corresponding StereoMapper cluster in which that entry was stored. This enabled direct comparison between the structure-based (StereoMapper) and context-based (MetaNetX) methods to assess how closely their resulting groupings aligned.

### 2.2 StereoMapper Workflow Overview

The following subsections describe each step of the StereoMapper workflow in detail, including structural processing and property extraction, the methodology underlying relationship prediction, and confidence score calculation. Together, these components provide a detailed overview of the pipeline’s operation and implementation. An overview of the pipeline is provided in Figure 1.

#### 2.2.1 Structure processing: standardisation and property extraction

Following data collection, all structures passed to StereoMapper were standardised and annotated with structural properties to prepare them for downstream analysis and ensure consistent comparison across sources. Standardisation of input structures is essential for accurate relationship assignment, as it enables consistent comparison of molecules originating from different databases and ensures the validity of resulting classifications. Challenges associated with representing organic structures have been described previously [29]. Consequently, converting all structures into a consistent format governed by a defined set of rules was a prerequisite for reliable comparison. To achieve this, a standardisation and property extraction procedure was applied to generate canonical representations and capture key structural features (e.g., formal charge, structural repeat unit presence). Two types of structural repeat unit are defined in this work: (i) *undefined structural repeat units,* cases where the repeating unit is not explicitly specified, resulting in a partially defined structure, and (ii) *defined structural repeat units,* cases where the repeating unit is fully specified, enabling explicit representation of the metabolites structure. Standardisation was performed using the method described by O’Boyle [30], implemented via the Open Babel cheminformatics package [18]. Two canonicalisation strategies are available; (i) Inchified SMILES and (ii) Universal SMILES. For this work, Inchified SMILES was selected, as it employs the InChI normalisation protocol that collapses recognised tautomers into a single representation. Consequently, recognised tautomeric variants are clustered as equivalent in this version of the pipeline, and a dedicated tautomer relationship is not assigned. This design prioritises cross-database harmonisation over explicit handling of tautomeric species. The procedure also assigns canonical atom labels, which are essential for accurate stereochemical comparison and atom-level mapping.

The accuracy of StereoMapper therefore depends on the robustness of the InChI software, reported to achieve >99.99% reliability [31], though minor limitations remain, particularly in the handling of some organometallic compounds. When canonicalisation was unsuccessful, a fallback procedure implemented with RDKit [17] ensured complete coverage by drawing on the canonical SMILES representation provided by RDKit. In parallel, several structural properties were extracted and recorded to guide downstream relationship assignment. These included formal charge, the first block of the InChIKey, and the presence of structural repeat units. Each of these properties are used downstream to generate isomeric sets and assign relationships. All processed structures and their associated features were stored in a dedicated structure cache database, ensuring that canonical identifiers and key properties were readily accessible for downstream analysis. These features guided decisions on whether structures were clustered as identical species or treated as distinct yet related molecules, requiring explicit relationship evaluation (see Section 2.2.2). As a result, all input structures were represented in a consistent canonical form and enriched with structural properties, providing a robust foundation for downstream clustering and relationship assignment within StereoMapper.

#### 2.2.2 Relationship Prediction Pipeline

With all structures standardised and annotated, StereoMapper performs three sequential operations: (i) clustering of equivalent metabolites to remove redundancy, (ii) prediction of stereochemically defined relationships between related molecules, and (iii) quantitative confidence scoring of each predicted relationship class. These stages collectively enable a stereo-aware, high precision mapping of molecular relationships.

##### Clustering and relationship assignment

Before predicting structural relationships, equivalent metabolites were clustered into single nodes, referred to as an isomeric set. Clustering offered two main advantages: (i) storage efficiency, by grouping equivalent structures into a single database entry, and (ii) computational efficiency, by avoiding unnecessary pairwise comparisons. Clustering was based on Inchified SMILES, formal charge, and structural repeat unit presence, as retrieved from the structure cache (see Supplementary Materials 2.1 for detailed schema). Structures were deemed identical if they contained the same atoms in the same spatial arrangement and shared the same protonation state. This definition captures molecule-level identity and intentionally excludes context-dependent effects such as pH. Candidate clusters were first defined by strict matching of Inchified SMILES and InChIKeys. Structural repeat unit logic (see Section 2.2.1 for definitions) was then applied. Undefined structural repeat units were clustered only with structures sharing identical InChIKey, Inchified SMILES, and structural repeat unit property. Defined structural repeat units were clustered with structures matching both identifiers and repeat count, or with non-sructural repeat unit structures sharing both identifiers. Each unique cluster was assigned a representative numeric identifier for traceability. Cluster member database identifiers were also recorded, extracted primarily from structure-data file (SDF) properties or, where unavailable, from molfile basenames. The cluster identifier and first InChIKey block of each cluster were then used to group related structures for relationship prediction. Clusters sharing the same first InChIKey block were compared, and for each cluster, the Inchified SMILES and associated features were retrieved.

To enable stereo-aware relationship assignment, an RDKit based procedure was implemented to extract stereochemical descriptors, including tetrahedral and cis/trans configurations. These descriptors were integrated into the structural comparison step, which also considers previously extracted molecular features such as charge and structural repeat unit presence. The comparison function combined these structural and stereochemical features to perform a stereochemically informed similarity assessment. Relationship classes were assigned according to stereochemical and structural correspondence rules consistent with those defined in Section 2.2.1. Two additional relationship types were introduced to capture cases where StereoMapper could not assign a definitive or reliable classification. The *unclassified* relationship indicates scenarios where StereoMapper recognises a structural relationship between molecules but the case is too complex to fit into an existing category. The *unresolved* relationship, on the other hand, denotes instances where the pipeline may have incorrectly identified two structures as equivalent. Such cases represent pipeline errors, as all truly equivalent structures are clustered in the preceding step. These occurrences are rare and are hypothesised to result from occasional stereochemical handling issues within RDKit. In cases where the primary approach failed—for example, due to structure file parsing errors or other processing issues—a fallback method based on InChI string matching was applied. This fallback mechanism maximised coverage while minimising errors arising from cheminformatics library limitations or malformed structure files. Use of the fallback method was explicitly reported to the user, both in the generated log files and in the extra info column of the relationships table within the output database (see Supplementary Materials, 2.2 for detailed schema). The resulting output is subsequently processed to assign a relationship type and a quantitative confidence score for each analysed structure pair.

##### Confidence Score Calculation

To quantify the reliability of each predicted relationship, a class-specific confidence score was computed. This score reflects how strongly the observed structural and stereochemical patterns agree with those expected for the assigned relationship class (e.g., protomers, enantiomers, diastereomers, and stereoresolution pairs). The calculation proceeds in three stages, each contributing additional layers of confidence: (i) *Stereo-consistency multiplier* (*M_stereo_*) — a class-specific multiplier computed from the stereochemical agreement between two structures. It accounts for the total number of stereogenic features, tetrahedral inversions or matches, double-bond inversions or matches, and undefined stereocentres. This term is normalised between 0 (no consistency) and 1 (maximum consistency); (ii) *geometric and topological similarity* — structural similarity captured using the normalised RMSD (*R_RMSD_*) of atom pairwise distances and the normalised 2D Tanimoto coefficient (*T*_2_*_D_*) derived from molecular fingerprints, both calculated with RDKit; and (iii) *final composite score* (*S*) — all components were integrated according to

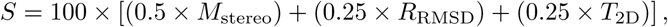

where *M*_stereo_ is the stereo consistency multiplier, *R*_RMSD_ is the normalised RMSD, and *T*_2D_ is the Tanimoto similarity respectively. Scores range from 0 to 100, with higher scores indicating stronger consistency across evaluated dimensions. High confidence classifications (*≥* 90) were considered reliable, while medium (70-89), low (40-69) and very low (*<* 40) scores signal relationships that may require manual review.

### 2.3 Benchmarking Experiments

This section describes the benchmarking framework used to evaluate the performance of StereoMapper in reproducing and extending stereochemical and structural relationships.

#### 2.3.1 Stereochemical Relationship Assignment

Benchmarking was conducted using all control datasets described in Section 2.1.1. StereoMapper was applied to each dataset to generate a set of predicted relationships. Because the pipeline checks for possible isomeric relationships, unique pair keys were used to filter out extra relationships and prevent discrepancies in accuracy calculations. For each relationship class *c*, evaluation was restricted to the curated set of positive pairs *G_c_*. Off-pair predictions (i.e., predictions on pairs not in *G_c_*) were excluded as out-of-scope. Within *G_c_*, incorrect class assignments were counted as false negatives for the true class and false positives for the predicted class. The resulting precision, recall, and F_1_ scores therefore represent in-scope, pair-scoped metrics. Some false negatives were observed, arising from (i) clustering of related species into a single node by StereoMapper, (ii) structural file parsing errors, or (iii) incomplete exclusion of structures containing wildcard atoms. In the case of clustering, this could occur either correctly or incorrectly depending on the structural relationships present. Since the control datasets consisted only of positive examples, true negatives could not be defined. Accordingly, performance was evaluated using precision, recall, and the F_1_ score, defined as

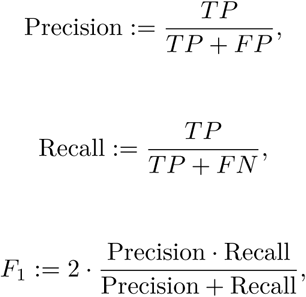

where true positives (TP) are predictions matching the control label, false positives (FP) are mismatches, and false negatives (FN) are pairs for which no classification could be assigned due to complex or ambiguous relationships.

#### 2.3.2 Comparison with existing mappings in a context-based framework

To evaluate the concordance between mappings generated by StereoMapper and those in the human subset of MetaNetX (derived from MNXref [32]), StereoMapper clusters were compared with the corresponding MetaNetX clusters as detailed in Section 2.1.2. As the two approaches apply different definitions of metabolite equivalence, dedicated agreement metrics were employed (see Figure 6). A summary of how each approach defines molecular equivalence is provided in Table 1. MetaNetX typically groups structurally related molecular species of a metabolite, such as protomers and stereoisomers, under a single identifier, combining structural information with reaction context. StereoMapper, in contrast, applies a structure-focused definition based on atomic connectivity, spatial configuration, and protonation state. As a result, a MetaNetX cluster generally contains more identifiers than a corresponding StereoMapper cluster. A follow-up analysis was performed to explore the cases in which the two approaches produced different groupings. For each such cluster, all relationship types assigned by StereoMapper were retrieved to generate a cluster-weighted distribution of structural relationships. It was anticipated that most differences would be associated with protomeric relationships, reflecting the differing equivalence definitions used by the two approaches.

**Table 1:** Comparison between StereoMapper and a context-aware framework (MetaNetX).

| Category | StereoMapper | Context-aware (MetaNetX) |
| --- | --- | --- |
| Basis | Purely structure-based; no contextual information used. | Integrates reaction co-occurrence with structural information within a reconciliation workflow. |
| Equivalence | Assigned when structures show matching molecular skeleton, spatial configuration and protonation. | May group structural variants together when reaction context supports equivalence. |
| Relationship assignment | Distinguishes multiple structural relationship types at isomer level. | Primarily defines parent-child relationships, often described broadly. |
| Protomer handling | Treats protomers as distinct entities, enabling recognition of protonation-dependent variants. | Often consolidates protonation states under a single identifier. |
| Stereochemistry | Explicitly distinguishes stereochemical relationships (e.g., enantiomers, diastereomers). | Stereochemistry often handled implicitly; contextual factors may cause consolidation under one identifier. |
| R-group / wildcard atoms | Excluded from analysis. | May be represented with contextual information, though non-trivial to interpret. |
| Software dependencies | RDKit, Open Babel, and InChI. | InChI, RDKit, and in-house Perl code. |
| Output type | SQLite database. | TSV and RDF files. |
| Limitations | Does not process R-groups; limited handling of complex relationships; focused on small molecules. | Limited relationship assignments, focused primarily on small molecules |

### 2.4 Data analysis and visualisation

Descriptive analyses of StereoMapper outputs, including relationship type distributions and cluster composition, were performed directly on the results generated by StereoMapper. These analyses were used to summarise and interpret StereoMapper performance when applied to the set of human structures extracted from MetaNetX.

## 3 Results

### 3.1 Benchmarking Relationship Assignment

The performance of StereoMapper was assessed using curated control datasets (see Section 2.1.1), where each dataset represents a distinct class of stereochemical or structural relationship (enantiomers, diastereomers, protomers, and stereo-resolution pairs), allowing comprehensive assessment. Precision, recall, and F_1_ scores were calculated for each dataset, and discrepancies were manually examined to distinguish genuine misclassifications from inconsistencies in the reference data. Across all datasets, StereoMapper demonstrated consistently high agreement with reference relationships (Figure 2). In the enantiomer set, most discrepancies arose from database inconsistencies rather than misclassification by StereoMapper, with several cases revealing previously unrecognised diastereomeric relationships in ChEBI. After accounting for these annotation mismatches, performance improved marginally, confirming strong reliability in distinguishing mirror-image structures. An example of StereoMapper-ChEBI disagreement is shown in Figure 8 (A). For diastereomers, StereoMapper maintained high accuracy, though minor deviations from estimated precision reflected the inherent difficulty of edge cases involving mixed stereochemical features. In the protomer dataset, false positives were primarily due to complex or ambiguous relationships that fall beyond the current scope of the pipeline, while most false negatives stemmed from clustering behaviour that merged related forms. After correction, performance approached the upper limits achievable under ideal conditions, with non-curated results reflecting more realistic operational performance. In the stereo-resolution pair dataset, StereoMapper again showed strong agreement, with most false positives linked to highly complex stereochemical scenarios and only a handful representing genuine misclassifications. Once adjusted, accuracy and consistency across this class were near complete, underscoring the robustness of the pipeline for challenging stereochemical distinctions. Overall, StereoMapper demonstrated high precision and recall across all relationship classes, performing best for enantiomeric and protomeric comparisons, and maintaining strong accuracy even for the more intricate stereo-resolution relationships. These results collectively highlight StereoMapper’s reliability in resolving fine-grained stereochemical relationships and its robustness under both controlled and real-world conditions.

**Figure 2:**
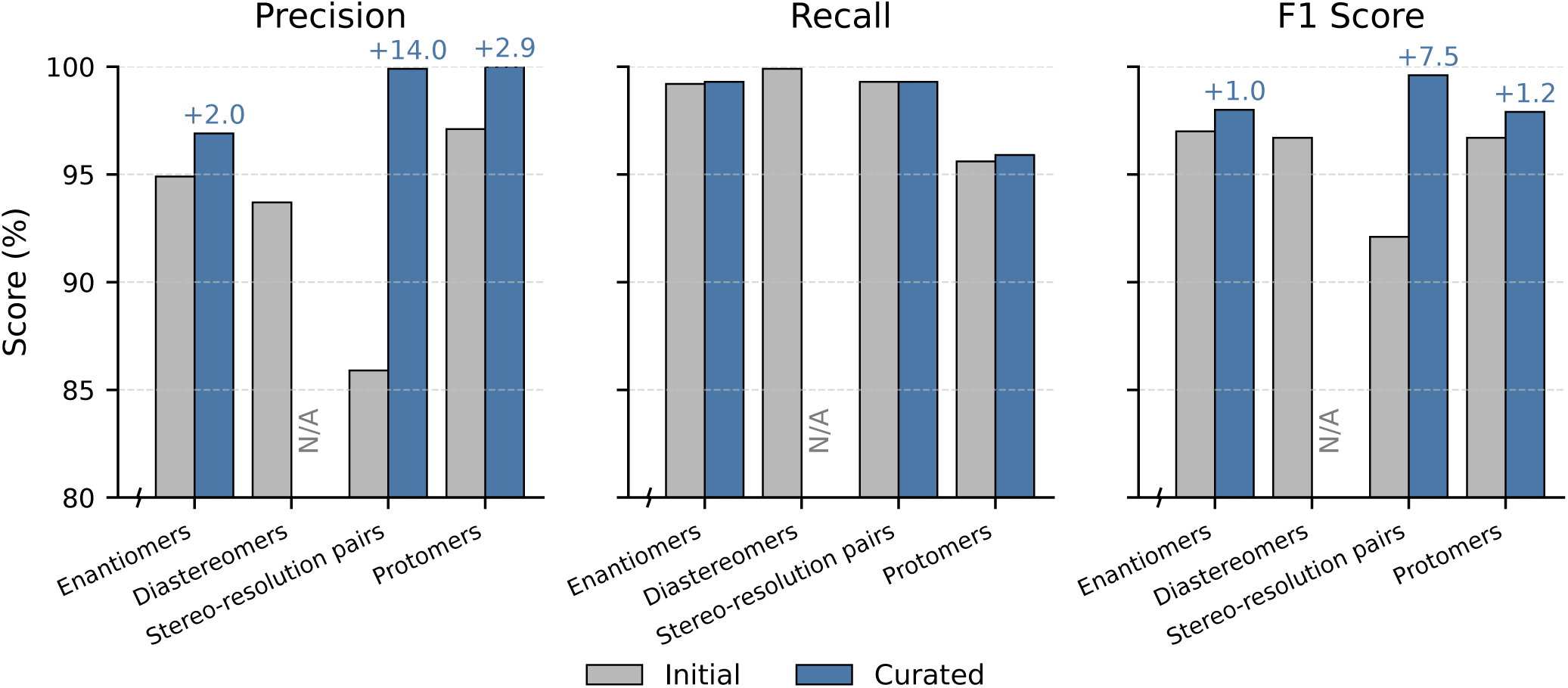
Benchmarking Performance by Relationship Type (initial vs curated). Each panel presents one evaluation metric (Precision, Recall, or F_1_ score), with the y-axis restricted to 80–100 percent as no variation occurred below this range. For each relationship class (Enantiomers, Diastereomers, Stereo-resolution pairs, and Protomers), grey bars indicate initial performance and blue bars indicate curated performance. Initial values correspond to the results obtained directly from the first pipeline outputs on each benchmark dataset. Curated values reflect cases in which false positives were manually examined to identify the underlying cause; where a putative error was found to be a correct assignment by StereoMapper, the performance score increased accordingly. Numerical labels above the blue bars show the change (Δ) relative to the initial score. Curated values are not reported for Diastereomers, as manual curation was not applicable. Overall, Precision and F_1_ score improved following curation, whereas Recall remained unchanged.

### 3.2 Application to the human subset of MetaNetX

#### 3.2.1 Overview of relationship assignments

The StereoMapper pipeline was applied to the set of human structures in MetaNetX, comprising approximately 1.3 million molecular structures. A total of 202,534 structures containing wildcard atoms were identified and excluded from the analysis. The remaining ∼1.1 million fully defined structures formed the basis for isomeric set and relationship generation. Because wildcard-containing entries were excluded, reported concordance values in Figure 6 apply only to fully defined molecules. Given that these excluded entries accounted for approximately 15% of the dataset, the estimates are representative of explicit, structure-defined compounds. StereoMapper produced 339,458 pairwise relationships. The distribution of assigned relationship classes is shown in Figure 3. Diastereomeric and enantiomeric relationships were the most frequently observed, whereas protomeric and unresolved cases were comparatively rare. This distribution provides a global overview of how StereoMapper assigns stereochemical relationships across the human content of MetaNetX. The distribution of StereoMapper confidence levels across relationship classes is shown in Figure 4. Unclassified and unresolved cases were omitted as they do not carry confidence scores. Among the remaining classified relationships (n = 311,593), the great majority were assigned with high confidence, with only a small proportion receiving low confidence.

**Figure 3:**
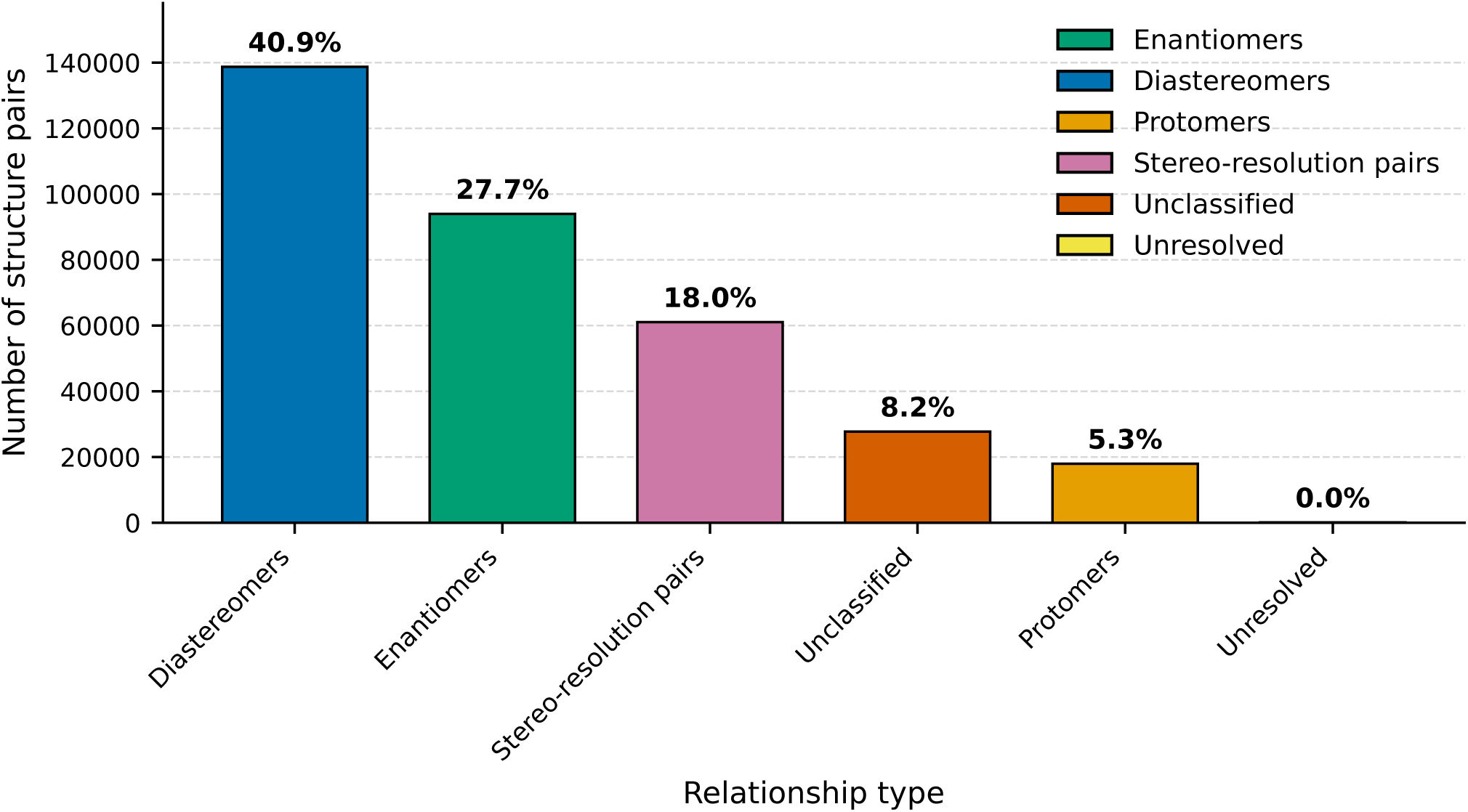
Distribution of StereoMapper relationship assignments across human MetaNetX structures (n = 1.3 million). Each bar represents a distinct stereochemical or structural relationship identified between molecular structure pairs by StereoMapper when applied to the set of human MetaNetX structures used in this study. Colours correspond to relationship types as indicated in the legend and are used consistently throughout the manuscript. A total of 339,458 relationships were identified, with diastereomers and enantiomers being the most frequently assigned classes.

**Figure 4:**
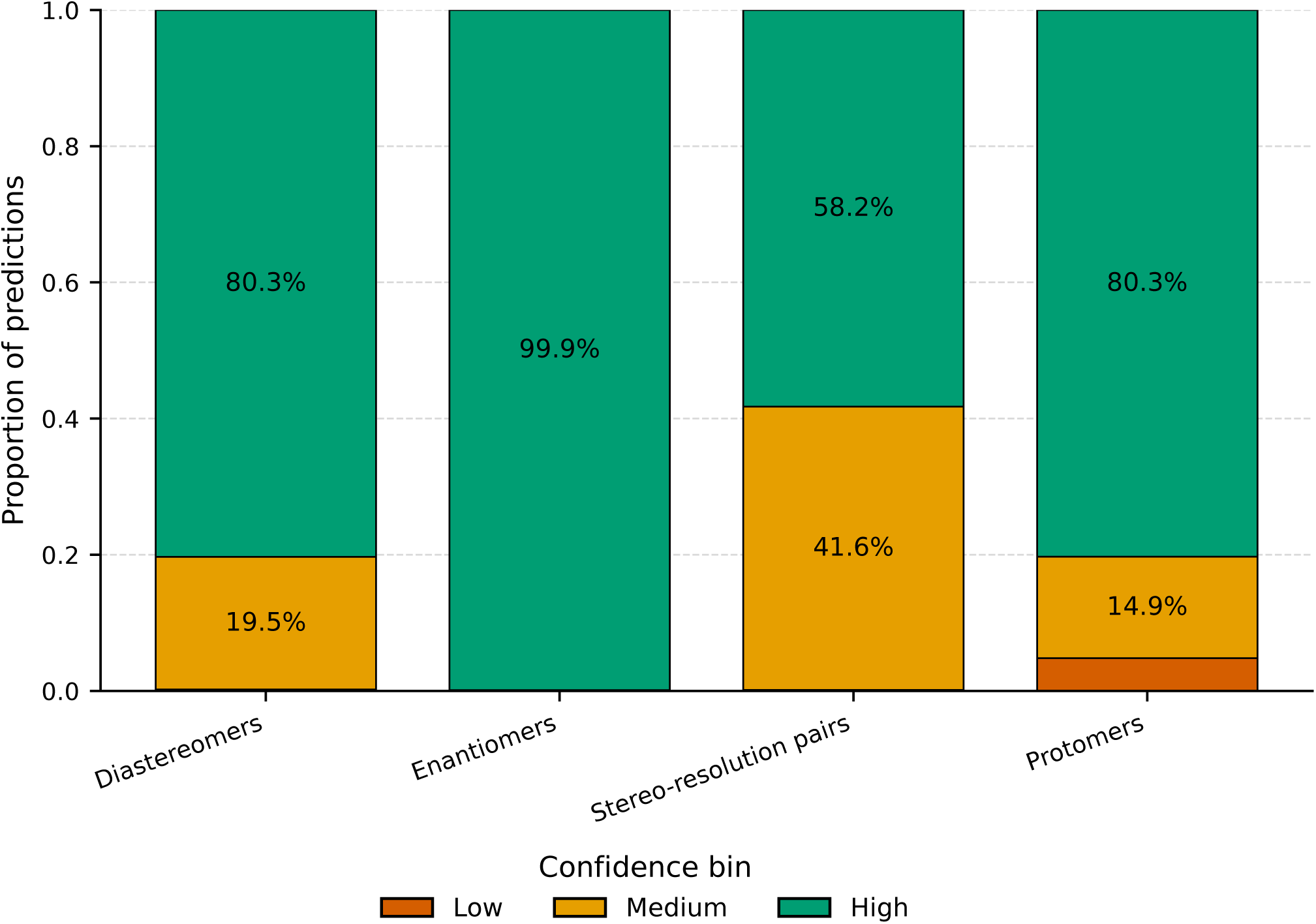
Distribution of StereoMapper confidence levels across relationship classes. Bars show the relative proportion of predictions assigned to each confidence level (Low, Medium, High) within every relationship class. The very low confidence level was not included as no relationship fell into this level. Unclassified and unresolved cases were excluded. This visualisation highlights the predominance of High-confidence assignments across most classes, indicating strong internal consistency of StereoMapper’s confidence scoring.

#### 3.2.2 Cluster composition

The identity clustering step produced a strongly right-skewed distribution of cluster sizes (see Figure 5). Most clusters consisted of a single structure (92%), while multi-member clusters were comparatively rare, with size-2 and size-3 clusters accounting for 5.4 % and 1.5 % of the total, respectively. Larger clusters (>3 members) occurred infrequently, indicating that groups containing more than three structures are uncommon at the StereoMapper identity level. Overall, this distribution indicates that most structures from the human portion of MetaNetX are unique when evaluated using StereoMapper.

**Figure 5:**
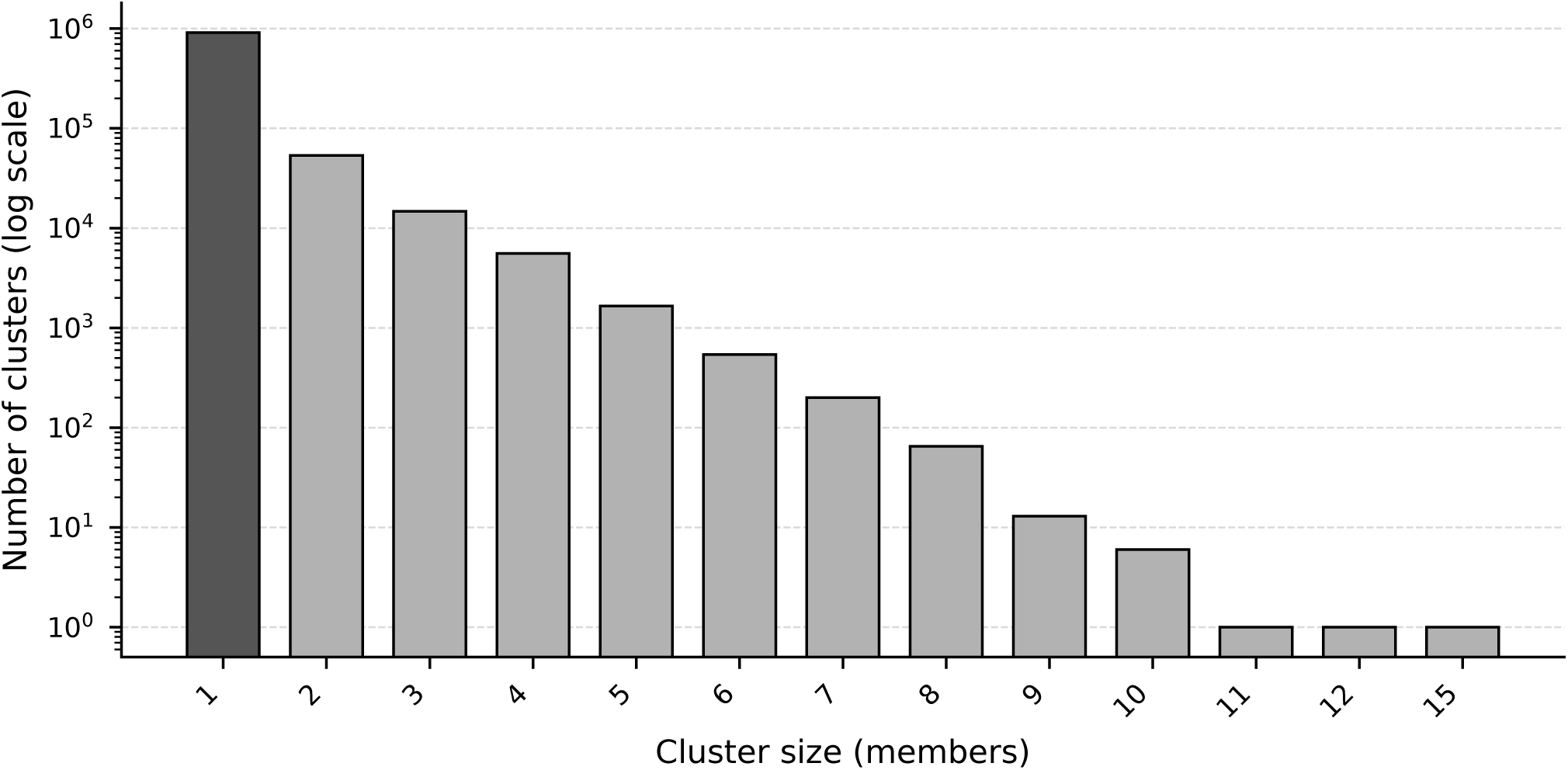
Distribution of StereoMapper isomeric set sizes on structures from representative databases (log-scaled y-axis). Most clusters are singletons (92%; bar shown in black), with the frequency of larger clusters decreasing exponentially. Bars are otherwise shown in uniform grey for clarity. The procedure used to generate identity clusters is described in Section 2.2.2.

#### 3.2.3 Comparison with existing mappings in MetaNetX

The majority of mappings between StereoMapper and the human subset of MetaNetX were consistent (total = 93.5%; green wedges in Figure 6). A smaller subset (6.5%; n = 63,774), showed non-matching groupings, reflecting instances where the two approaches applied different definitions of molecular equivalence. To explore the nature of these divergent groupings, the relationship classes represented within this subset (red wedge) were examined using the accompanying breakdown chart, as indicated by the grey dashed connection. Most differences were associated with protomeric variation (52.3%), followed by stereo-resolution relationships (37.4%). Unclassified cases contributed 21.6%, while diastereomeric and enantiomeric distinctions appeared rarely. These findings indicate that most non-matching groupings arise from differences in protonation handling or from situations in which the underlying resources provide structures with varying degrees of stereochemical specification. The remaining cases reflect complex or ambiguous structures that fall outside the current scope of StereoMapper. Overall, StereoMapper and MetaNetX show a high degree of consistency, with only a small proportion of cases yielding different equivalence groupings (Figure 6).

**Figure 6:**
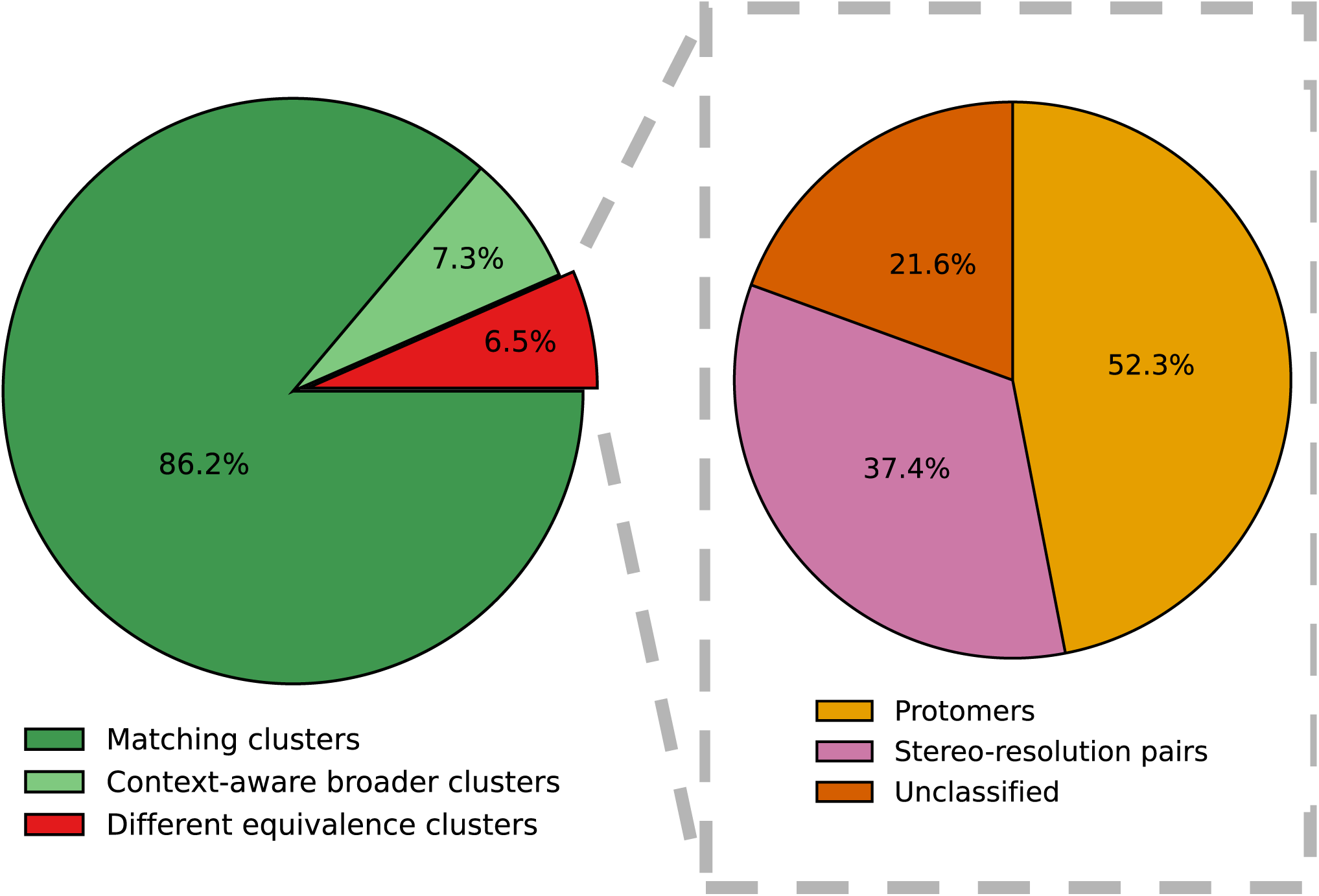
Mapping Concordance Between StereoMapper and human-MetaNetX, and the Structural Relationships Present Within Disagreement Cases. The pie chart on the left summarises mapping concordance between the structure-based (StereoMapper) and context-aware (MetaNetX) approaches. Green wedges represent cases where the two methods show consistency in structural groupings, whereas the red wedge denotes cases in which the two methods showed differences in groupings. The legend beneath the chart specifies the three concordance categories: matching clusters, context-aware broader clusters, and different equivalence clusters. *Matching clusters* corresponds to cases where the identifiers contained within clusters from both methods match exactly. *Context-aware broader clusters* describes cases where the mappings agree, but the MetaNetX cluster contains additional identifiers, typically arising from secondary accessions. *Different equivalence clusters* refers to cases where the clusters produced by the two approaches do not align. The pie chart on the right shows the distribution of structural relationship types identified within the different equivalence clusters set. Colours follow the scheme used in Figure 3. The wedges do not sum to one hundred per cent, as this analysis reports the proportion of disagreement clusters in which each relationship type occurs at least once. Rare classes, including enantiomers and diastereomers, are not shown. The legend beneath the chart identifies the three relationship classes depicted: protomers, stereo-resolution pairs, and unclassified. The grey dashed lines stemming from the different equivalence clusters wedge (red, left pie chart) encapsulating the pie chart on the right, indicate that the pie chart on the right are the distributions of relationships assigned within the strict disagreement cluster.

An example where StereoMapper and MetaNetX diverge is shown in Figure 7. The figure illustrates a case in which StereoMapper distinguishes several structural variants that MetaNetX treats as functionally equivalent within its context-aware definition of identity. This illustrates how the two approaches capture different dimensions of identity. StereoMapper separates these variants into four clusters and assigns relationship classes between them. This illustrates how structural and context-derived mappings can together offer a clearer and more chemically grounded view of metabolite variation.

**Figure 7:**
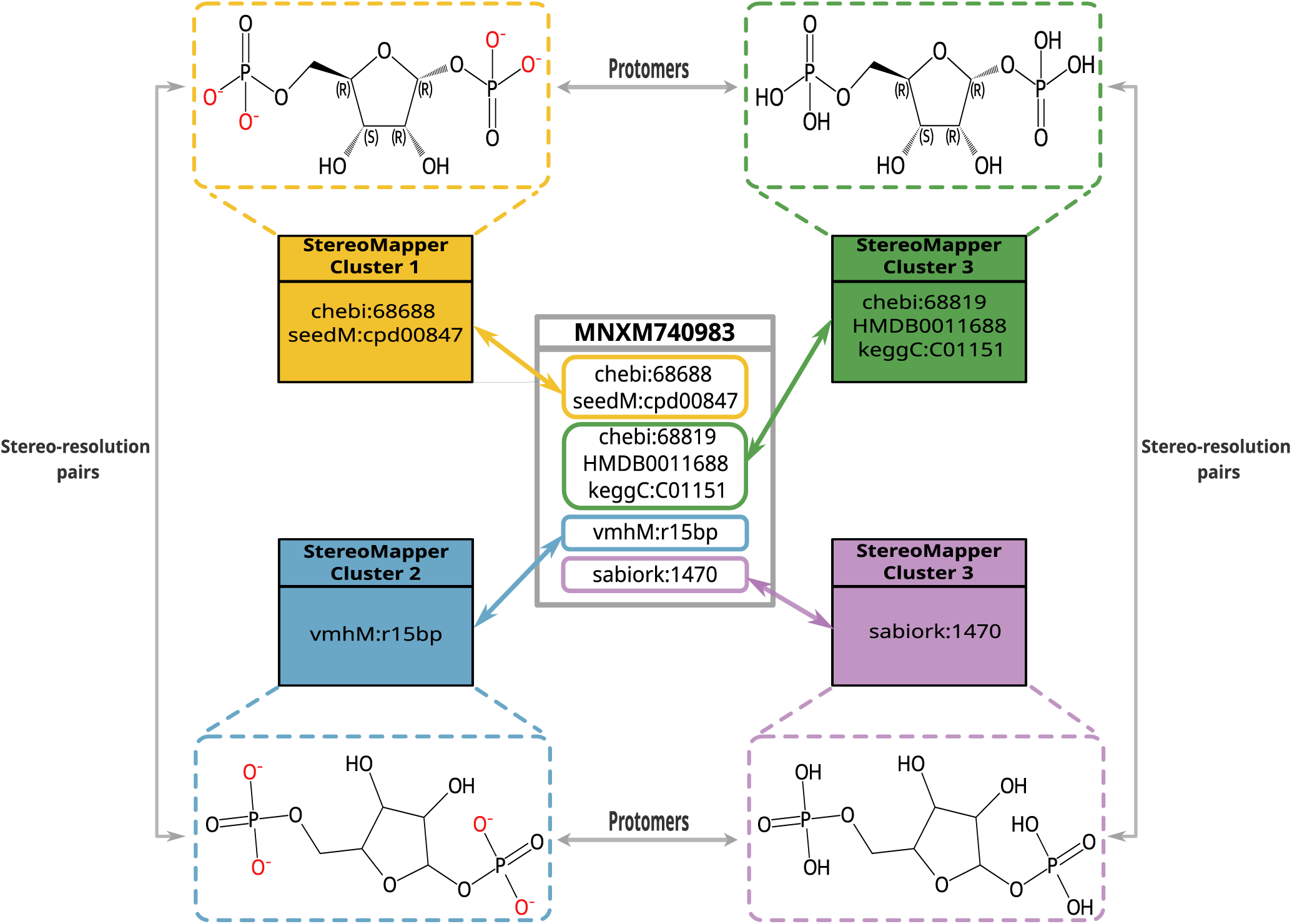
Structural variation within a context-aware identifier illustrated by StereoMapper clustering. Visual summary of a cluster in which a single identifier within a context-aware framework (MetaNetX) encompasses several metabolite structures that differ at the level of fine-grained stereochemical or protonation detail. Colour-coded boxes illustrate how StereoMapper differentiates these structures into separate, structure-defined clusters. Four such clusters are shown for this MetaNetX identifier, each displayed within a colour-coded dashed grid. Grey bidirectional lines between grids depict the structural relationship classes inferred by StereoMapper. Together, these elements demonstrate how a structure-based perspective can complement a context-aware framework by providing additional resolution on the structural variation present within an identifier.

### 3.3 Illustrative examples of StereoMapper’s classification improvements

StereoMapper demonstrates its ability to identify, correct and enhance relationship classifications within the reference databases it interacts with. During benchmark testing, for instance, StereoMapper correctly reclassified a misassigned relationship in ChEBI, identifying a diastereomeric pair where the database had incorrectly recorded an enantiomeric one. This case is shown in Figure 8 (A). Beyond correcting existing misclassifications, StereoMapper also enriches current relationship types. In Figure 8 (B), a parent-child relationship recorded in MetaNetX is refined by StereoMapper to an explicit diastereomeric assignment. In some cases, the structural perspective provided by StereoMapper offers an alternative interpretation that complements the context-aware grouping, as seen in Figure 8 (C). This example, which also forms part of Figure 7, shows two structures that a context-aware framework treats as functionally equivalent. Alternatively, StereoMapper identifies them as distinct metabolite variants and assigns them to two separate clusters with a protomeric relationship. These examples represent only a small subset of the many cases observed in this study, and together highlight the complementary role that a structure based pipeline can play when applied alongside a context aware framework.

**Figure 8:**
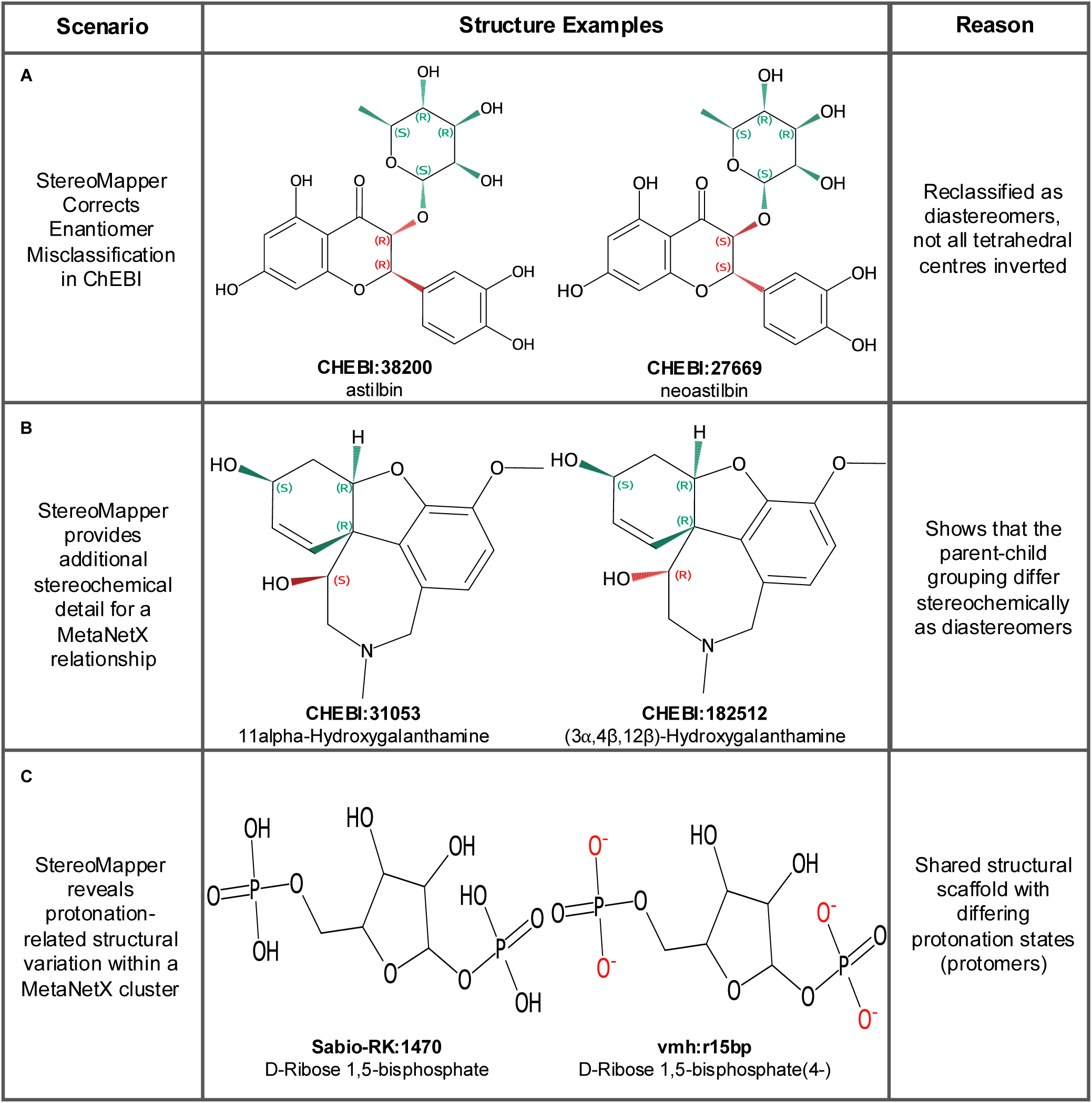
Examples of StereoMapper performance. Each row illustrates a distinct use case, showing the scenario, corresponding structural examples, and the rationale for assignment (as indicated by the column headers). (A) StereoMapper corrects a misclassification in ChEBI, reassigning a pair previously labelled as enantiomers to the diastereomer class. (B) StereoMapper provides additional stereochemical detail for an existing parent–child relationship in MetaNetX by specifying the relationship class (diastereomer). (C) StereoMapper reveals a protonation related structural variation within a single MetaNetX cluster, correctly labelling them as protomers. Together, these examples demonstrate the versatility of StereoMapper in improving curation accuracy and facilitating reliable cross-mapping.

## 4 Discussion

### 4.1 Background

Context-aware resources such as MetaNetX [11] define metabolite equivalence based on reaction co-occurrence with some integration of structural information, which can lead to broader groupings than a purely structural perspective. StereoMapper provides a complementary, structure-centred view by resolving stereochemical and protonation differences that may be obscured in context-aware mappings. This framing guides the interpretation of the results, particularly in cases where differences arise in how the two approaches organise metabolites.

### 4.2 Evaluation and interpretation of StereoMapper

#### 4.2.1 Benchmarking relationship assignment

StereoMapper demonstrated strong performance across all benchmarking tests (Figure 2). All relationship classes exhibited high precision, recall, and *F*_1_ scores, indicating that StereoMapper can accurately assign relationships across all defined classes. The predominance of high confidence scores across each relationship class reinforces that the majority of relationship assignments are highly likely to be true. A key observation concerns the comparison between initial and curated results: for all relationship types (except diastereomers; see 2.1.1), both precision and *F*_1_ score improved following adjustment. These two evaluations represent distinct scenarios: (i) real-world conditions, where database inconsistencies persist, and (ii) an optimal scenario, where StereoMapper identifies and compensates for such errors. These findings suggest that StereoMapper not only achieves high overall accuracy, but also possesses the capability to detect potential inconsistencies in existing databases (Figure 8 A). This demonstrates the unique capability of StereoMapper to both establish new relationships and refine existing ones. Taken together, the benchmarking results confirm that StereoMapper provides reliable, stereochemically consistent relationship assignments between related structures, reinforcing its role in closing the relationship gaps identified earlier.

#### 4.2.2 Relationship analysis and cluster composition

Interpreting the outputs of **StereoMapper** when applied to the human content of MetaNetX provides insight into how the pipeline builds on existing mappings by defining new structural relationships within the context-aware framework.

##### Relationship analysis

The predominance of stereochemical relationships identified by StereoMapper (Figure 3) shows that most newly resolved associations stem from differences in spatial configuration rather than composition or charge. This confirms both that StereoMapper primarily contributes stereochemically informed distinctions and that many metabolites in the human MetaNetX set differ chiefly through stereochemistry. Together, these findings indicate that StereoMapper offers a complementary perspective by adding a stereochemistry-sensitive layer of connectivity to existing context-aware mappings, thereby improving the resolution and interpretability of structural associations. By defining these relationships explicitly, StereoMapper enhances the granularity of structural connectivity within a context-aware framework and provides curators and developers with richer information for relating compounds.

##### Cluster composition

Analysis of cluster composition provides a complementary perspective on how structural organisation is altered following StereoMapper integration. The vast majority of clusters generated by StereoMapper are singletons (Figure 5), indicating that the human portion of MetaNetX contains predominantly unique molecular structures. Larger clusters (*≥*3 members) are rare, suggesting that most structures are chemically distinct and that duplicate or near identical entries are uncommon. This implies that the integrated databases contribute diverse metabolic structural data, underscoring the need for harmonisation and reconciliation to improve GEM development. These findings reinforce the claim that StereoMapper maintains a high level of structural specificity, distinguishing truly equivalent compounds, while avoiding over-merging of related but non-equivalent structures. Consequently, the observed cluster composition supports StereoMapper’s role in refining equivalence definitions. As a result, curators can confidently treat most of the structures within the human portion of MetaNetX as distinct molecular entities, when processed by StereoMapper, possibly simplifying manual curation tasks. Taken together, the results indicate that StereoMapper offers an additional structural perspective that complements context-aware reconciliation and supports clearer relational interpretation. The identification of more than 339,000 relationships, most of which reflect genuine structural distinctions, illustrates the benefit of adding stereochemical and protonation detail to existing mapping resources. In doing so, StereoMapper provides a complementary method, providing reliable molecular relationships, thereby supporting more accurate biocuration, reaction mapping, and downstream GEM reconstruction and prediction.

#### 4.2.3 Comparison with existing mappings from a context-aware framework

To evaluate the accuracy of the isomeric sets produced by StereoMapper, these sets were compared with the clusters defined in MetaNetX to assess the extent of concordance. In this setting, an isomeric set refers to a group of structures considered equivalent under the identity criteria used by StereoMapper, with MetaNetX clusters defined according to its own criteria, as shown in Table 1. It was anticipated that most groupings would overlap, with any differences reflecting the distinct principles each method applies when defining metabolite equivalence. It was also expected that groupings identified as diverging would be linked by relationships newly assigned by StereoMapper, illustrating how the structural perspective offers an alternative but complementary interpretation of these groupings.

##### Comparison of structural and context-aware cluster organisation

The comparison between StereoMapper groupings and the clusters defined in MetaNetX showed that most groupings aligned closely, with a smaller proportion falling into groupings defined differently by each method (Figure 6). This pattern is consistent with the expectations set out before analysis and reflects the distinct principles each approach uses when defining metabolite identity. These observations support the reliability of the structural groupings produced by StereoMapper and complement the patterns seen across structural composition classes (Figure 5). Overall, the comparison illustrates how a structure-centred view can operate alongside context-aware reconciliation to provide a more complete picture of metabolite relationships.

##### Sources of grouping differences

To understand the origins of the grouping differences, we examined the relationships assigned by StereoMapper within these sets. Protomers accounted for most of the distinct groupings observed (Figure 6), which aligns with the behaviour of context-aware frameworks that often place protonation variants within the same cluster, as summarised in Table 1. An example of such a case is provided in Figure 8 (C). Stereo-resolution pairs and unclassified cases followed in frequency, the latter reflecting complex or ambiguous structures that fall outside the current scope of StereoMapper. A recurrent pattern is that context-aware resources tend to define broad structural relationships, reflecting their focus on reaction-level equivalence. StereoMapper’s explicit relationship assignments offer a complementary, fine-grained view by providing additional chemical context. For instance, Figure 8 (B) shows a scenario in which StereoMapper refines a broad parent–child within MetaNetX grouping by indicating that the structures involved do not share a direct parent within the set of databases assessed. This illustrates how the structural perspective can enrich context-aware reconciliation by highlighting alternative ways in which compounds may be related. The added granularity also presents potential benefits for biocuration. Querying relationships generated by StereoMapper enables retrieval of all structural variants of a metabolite, offering two advantages for metabolic reconstruction: improved efficiency when identifying the most appropriate structural form, and the ability to generate alternative reactions involving these distinct variants. Enantiomeric and diastereomeric distinctions contributed only minimally to the observed grouping differences, indicating that these stereochemical features are already well represented in existing resources. Some stereochemical variation may, however, be embedded within the unclassified category, particularly where it coincides with differences in protonation state. Overall, the grouping differences primarily reflect the distinct principles each method applies, rather than conflicts in interpretation, highlighting how the two perspectives can operate together to provide a more complete representation of metabolite relationships.

Taken together, these findings show that the groupings produced by StereoMapper correspond closely with those in a well-established context-aware resource, while adding further interpretative depth through explicit structural relationship assignments. This additional detail supports clearer understanding of how chemically related entities connect and addresses earlier considerations regarding transparency in identity definitions. Rather than replacing context-aware frameworks, StereoMapper complements them by introducing a stereochemistry aware relational layer that enhances how molecular entities can be interpreted. Across the benchmarking analysis, relationship statistics, and comparison of grouping patterns, StereoMapper illustrates how a structural perspective can operate alongside existing approaches to provide a broader and more informative view of metabolite relationships.

The table outlines conceptual and technical differences between a structure-based pipeline (StereoMapper) an a context-aware framework (MetaNetX). Categories include the basis of the approach, defintion of equivalence, relationship assignment, stereochemical handling, protomer treatment, and data representation. The comparison highlights StereoMapper’s explicit structure-based approach versus the context-aware nature of MetaNetX, underscoring their complementary roles in metabolite harmonisation.

### 4.3 Implications, limitations and future perspectives

#### 4.3.1 Implications

StereoMapper provides a reliable pipeline for representing the structural dimension of metabolite equivalence by assigning explicit relationship classifications between variants of a compound, allowing structurally distinct forms to be represented separately where appropriate. In this study, its application to the human subset of a context-aware framework (MetaNetX) showed that StereoMapper adds structural precision and enriches relational information within the network. Beyond this case study, StereoMapper offers an additional resource for biocuration workflows. Curators can use its outputs to retrieve all structural variants of a given metabolite from the respective databases, supporting the identification of appropriate molecular forms during reaction annotation or model construction. It is worth noting that the StereoMapper pipeline is not limited to the databases examined here. Structures from any source can be used, provided they are correctly formatted and represent a defined chemical species. This stereochemically-aware search capability can substantially increase the efficiency and accuracy of manual curation, particularly in datasets where several structural representations of the same metabolite exist. At a broader scale, GEMs stand to benefit from these improvements. Greater clarity in metabolite cross-mapping can highlight structural mismatches or ambiguities within reaction definitions, improving both internal consistency and predictive reliability. More efficient and accurate biocuration further contributes to faster model development and higher data quality within the biochemical resources that underpin these models. Because it operates independently of database-specific conventions, StereoMapper can be readily integrated into other biochemical frameworks, offering a universal stereochemistry-aware layer for structural reconciliation across heterogeneous resources. In this way, StereoMapper provides a complementary approach to existing frameworks, extending relational context and clarifying structure-defined equivalence, thereby supporting enhanced reconciliation.

#### 4.3.2 Limitations and future directions

Although StereoMapper performs well across benchmarking and application tests, certain limitations define its current scope. The most immediate limitation is the reliance on having accurate structural representations of the metabolite of interest; if the supplied structure does not match the user’s intended compound, unexpected, though not incorrect, results will be produced. Partially defined structures containing R-groups are also excluded, as their undefined substructures preclude reliable stereochemical or equivalence comparisons. This remains an active area of research in cheminformatics [33, 34]. Additional constraints arise when compounds exhibit overlapping structural differences, such as protonation changes coinciding with stereochemical variation, which contributes to the unclassified cases described in Section 4.2.3, and Figures 3 and 6. Further refinement of StereoMapper’s rule set would be needed to capture these multidimensional relationships. Moreover, the present implementation has been validated primarily on small molecules relevant to metabolic modelling; extending its application to larger or polymeric species would require substantial algorithmic modification. A related limitation is the normalisation of tautomeric variants to single canonical forms, which prevents explicit assignment of tautomer relationships and represents a clear avenue for further development.

Several practical considerations also relate to implementation and reproducibility. The current pipeline depends on external cheminformatics libraries such as RDKit (2025.3.3), Open Babel (3.1.1), and InChI (1.07) [17, 18, 21]. Minor version differences in these toolkits can influence canonicalisation or stereochemistry perception, potentially affecting reproducibility across systems. To mitigate this, StereoMapper is distributed within a version-locked environment to ensure consistency across platforms. A further consideration is that both the diastereomer control set and the StereoMapper pipeline rely on RDKit for stereochemical interpretation, creating partial alignment between the stereochemical models used for curation and evaluation. This reflects a trade-off between independence and reproducibility, as RDKit currently offers the most comprehensive and transparent open-source support for stereochemical analysis. Nonetheless, future benchmarking using independently curated datasets and alternative toolkits would help to confirm generalisability and assess open-world specificity. A further potential limitation concerns the remaining benchmarking datasets, where the reliability of the relationships provided by ChEBI (for enantiomer and protomer sets) and MetaNetX (parent-child relationships) may affect the accuracy of the ground truth.

Looking ahead, several avenues for development are under consideration, with a central objective being the extension of StereoMapper beyond structure-level mapping towards reaction cross-mapping. This work supports the development of the ReconX knowledge graph (ReconXKG) [35], part of the Reconstruction and computational Modelling for Inherited Metabolic Diseases (Recon4IMD) project. ReconXKG aims to harmonise diverse biochemical resources, including those examined in this study, enabling the construction of hybrid GEMs that draw on all available evidence to improve model completeness and predictive accuracy. Such integration would help address current gaps in reaction-level overlap that limit reaction reconciliation within the graph. Future work will consider converting StereoMapper outputs into resource description framework (RDF) format for full incorporation into ReconXKG, as well as exploring integration of the StereoMapper pipeline within MetaNetX so that the two resources can operate in a complementary manner and offer users a more comprehensive form of reconciliation. Together, these efforts aim to transition StereoMapper from a standalone stereochemical analysis tool into a component of large-scale biochemical data harmonisation. The implications, limitations, and future directions outlined here illustrate a clear trajectory for the continued evolution of the pipeline. Addressing current constraints and expanding its scope would further establish StereoMapper as a valuable element within broader reconciliation workflows.

### 4.4 Conclusions

StereoMapper introduces a reliable, stereochemistry-aware pipeline for identifying and classifying relationships between related molecular structures. By distinguishing equivalent compounds from closely related variants, it provides structural clarity that is not explicitly captured in current cross-mapping resources. Benchmarking analyses, relationship statistics, and comparisons with the established context-aware MetaNetX dataset collectively demonstrate the robustness, interpretability, and interoperability of the StereoMapper approach. This work complements existing context-aware frameworks by adding a stereochemically informed layer of information that supports clearer identity reconciliation and a more detailed understanding of structural relationships. In doing so, StereoMapper contributes to more transparent biocuration and has the potential to strengthen the predictive reliability of genome-scale metabolic models.

## 5 Abbreviations

**Table 2:**
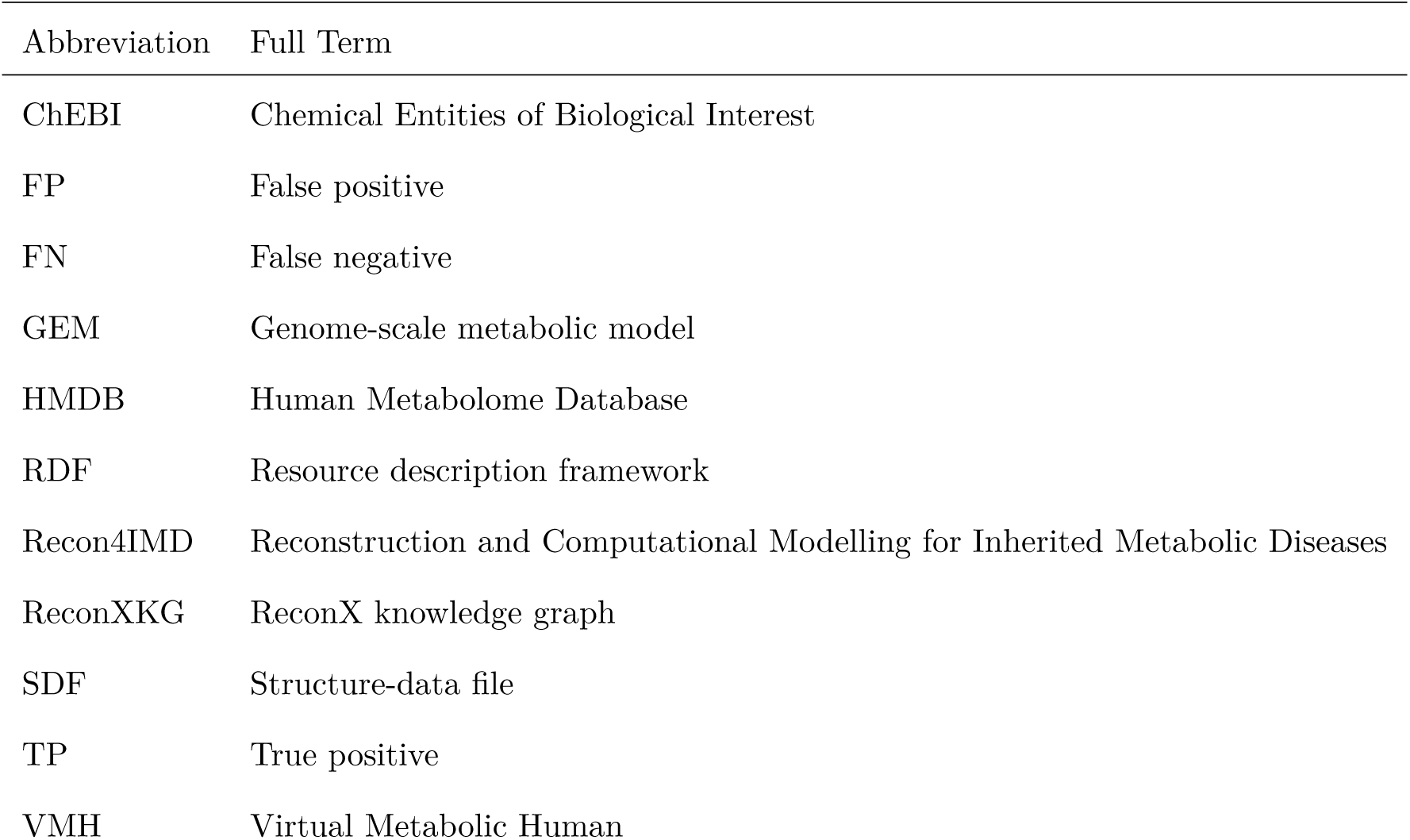
List of Abbreviations.

| Abbreviation | Full Term |
| --- | --- |
| ChEBI | Chemical Entities of Biological Interest |
| FP | False positive |
| FN | False negative |
| GEM | Genome-scale metabolic model |
| HMDB | Human Metabolome Database |
| RDF | Resource description framework |
| Recon4IMD | Reconstruction and Computational Modelling for Inherited Metabolic Diseases |
| ReconXKG | ReconX knowledge graph |
| SDF | Structure-data file |
| TP | True positive |
| VMH | Virtual Metabolic Human |

## 6 Declarations

### 6.1 Availability of data and materials

The StereoMapper pipeline is openly accessible in a public GitHub repository [36] which includes all source code, installation guidance, and the scripts used to generate the results reported here (available in the experiments directory). The databases containing the structures evaluated in this study are provided in the associated Zenodo repository [37]. Users wishing to retrieve VMH structures are directed to the VMH database [6]. The databases supporting the conclusions of this article are supplied in Additional Files 1 and 2. Structural representations are not distributed with the released databases, owing to the sensitive nature of a subset of structures used in this study. Consequently, some columns are omitted from the released database files, even though they appear in Section 2 of the Supplementary Materials, which describes the schema of the output databases.

### 6.2 Competing interests

The authors declare that they have no competing interests.

### 6.3 Funding

This research was supported by Recon4IMD, which is co-funded by the European Union’s Horizon Europe Framework Programme (101080997), the Swiss State Secretariat for Education, Research and Innovation (23.00232), and by United Kingdom Research and Innovation (10083717 & 10080153).

### 6.4 Authors Contributions

**Jack McGoldrick:** Methodology; Software; Formal analysis; Investigation; Data curation; Writing - Original Draft

**Marco Pagni:** Methodology; Conceptualisation; Writing - Review & Editing, Funding acquisition

**Saleh Alwer:** Software; Validation; Writing - Review & Editing

**Joanne Cooney; Natalia Makosa; Anne Niknejad; Sébastien Moretti; Jadzia Murphy; Filippo Martinelli**: Resources; Data curation

**Alan Bridge; Ines Thiele:** Resources, Data curation, Funding acquisition

**Ronan M. T. Fleming:** Conceptualisation; Methodology; Writing - Review & Editing; Supervision; Project administration; Funding acquisition

## Acknowledgements

We thank Anastasia Sveshnikova for her guidance in adopting a 2D stereochemically annotated workflow, which improved both the computational efficiency and accuracy of the pipeline.

## Notes

### Competing Interest Statement

The authors have declared no competing interest.

